# Genetic variants associated with motion sickness point to roles for inner ear development, neurological processes, and glucose homeostasis

**DOI:** 10.1101/002386

**Authors:** Bethann S. Hromatka, Joyce Y. Tung, Amy K. Kiefer, Chuong B. Do, David A. Hinds, Nicholas Eriksson

**Affiliations:** 23andMe, Inc., Mountain View, CA, USA

## Abstract

Roughly one in three individuals is highly susceptible to motion sickness and yet the underlying causes of this condition are not well understood. Despite high heritability, no associated genetic factors have been discovered to date. Here, we conducted the first genome-wide association study on motion sickness in 80,494 individuals from the 23andMe database who were surveyed about car sickness. Thirty-five single-nucleotide polymorphisms (SNPs) were associated with motion sickness at a genome-wide-significant level (*p* < 5 × 10^−8^). Many of these SNPs are near genes involved in balance, and eye, ear, and cranial development (e.g., *PVRL3*, *TSHZ1*, *MUTED*, *HOXB3*, *HOXD3*). Other SNPs may affect motion sickness through nearby genes with roles in the nervous system, glucose homeostasis, or hypoxia. We show that several of these SNPs display sex-specific effects, with as much as three times stronger effects in women. We searched for comorbid phenotypes with motion sickness, confirming associations with known comorbidities including migraines, postoperative nausea and vomiting (PONV), vertigo, and morning sickness, and observing new associations with altitude sickness and many gastrointestinal conditions. We also show that two of these related phenotypes (PONV and migraines) share underlying genetic factors with motion sickness. These results point to the importance of the nervous system in motion sickness and suggest a role for glucose levels in motion-induced nausea and vomiting, a finding that may provide insight into other nausea-related phenotypes such as PONV. They also highlight personal characteristics (e.g., being a poor sleeper) that correlate with motion sickness, findings that could help identify risk factors or treatments.

## Introduction

Motion sickness is provoked by exposure to a variety of motions (e.g., traveling in cars, boats, or planes; amusement park rides; skiing; and riding on camels) [1]. Simulators and virtual reality environments can also induce motion sickness [2]. Symptoms of motion sickness include dizziness, nausea, vomiting, headache, and pallor [3]. Sweating, drowsiness, increased salivation, hyperventilation, and emotional distress may also occur. Motion sickness is associated with other conditions including migraines, vertigo, postoperative nausea and vomiting (PONV), and chemotherapy-induced nausea and vomiting (CINV) [1, 4].

Roughly one in three individuals is highly susceptible to motion sickness and the rest of the population may experience motion sickness under extreme conditions [5]. The underlying etiology of motion sickness, however, is not well understood. One theory suggests that motion sickness results from contradictory information the brain receives during motion [1, 5]. The vestibular system of the inner ear, which senses motion and body position and influences balance, signals “moving” to the brain, while the eye signals “stationary” because the car or boat appears stationary relative to the viewer. The vestibular system is also thought to serve as a sensor of disequilibrium-causing neurotoxins (i.e., a toxin detector) and is believed to trigger the emetic response in order to rid the body of toxins. Thus, motion sickness may be an aberrant trigger of the emetic response. Evidence for the involvement of the vestibular system comes from the observation that individuals with complete loss of the vestibular apparatus, a component of the vestibular system, are immune to motion sickness [1].

A variety of factors influence risk for motion sickness. Women are more susceptible than men [6–9] and younger individuals are at increased risk [8, 9]. Ancestry may also play a role; there is some evidence that motion sickness occurs more frequently in individuals with Asian ancestry compared to European ancestry [10, 11]. Some variables are situational and/or behavioral. For instance, one study showed that passengers without a view of the road ahead were about three times more likely to experience illness [8] and another report suggested that adopting a wider stance may reduce motion sickness [12]. There is also evidence that diet and eating behavior influence risk [7].

Perhaps the most important and least understood variable is the underlying physiological susceptibility of the individual. In women, increased cortisol levels are predictive of motion sickness [13] and susceptibility to motion sickness changes as a function of the menstrual cycle, suggesting that levels of estrogen and other hormones might play a role [14]. In both sexes, hyperglycemia is implicated in motion-induced nausea and vomiting [15]. There is also some evidence that lower baseline levels of adrenocorticotropic hormone (ACTH) [16], also known as corticotropin, and low sympathetic nervous system activity [17] increases susceptibility. Finally, since antihistamines (e.g., Dramamine), anticholinergics (e.g., scopolamine), and sympathomimetics (e.g., d-amphetamine and ephedrine) are effective treatments, altered baseline activity of the receptors these drugs bind to might influence risk for motion sickness.

Although heritability estimates for motion sickness range from 57–70% [18], genome-wide association studies (GWAS) on this phenotype have not been reported. Here, we describe a large GWAS in which we find 35 regions significantly associated with motion sickness.

## Results

### Genome-wide Association Study of Motion Sickness

We performed a GWAS in 80,494 individuals from the customer base of 23andMe, Inc., a personal genetics company. Participants were of primarily European ancestry and were at most distantly related to each other (i.e., first cousins and closer were excluded). Motion sickness was assessed using online self-report. Participants responded to questions about their degree of car sickness and questions were combined onto a scale of 0 (never motion sick), 1 (occasionally), 2 (sometimes), or 3 (frequently). Details about the cohort can be found in Table 1 and in the Methods. All analyses were controlled for age, sex, and five principal components of genetic ancestry. Manhattan and quantile-quantile plots are provided in Figures 1 and S1.

**Figure 1:**
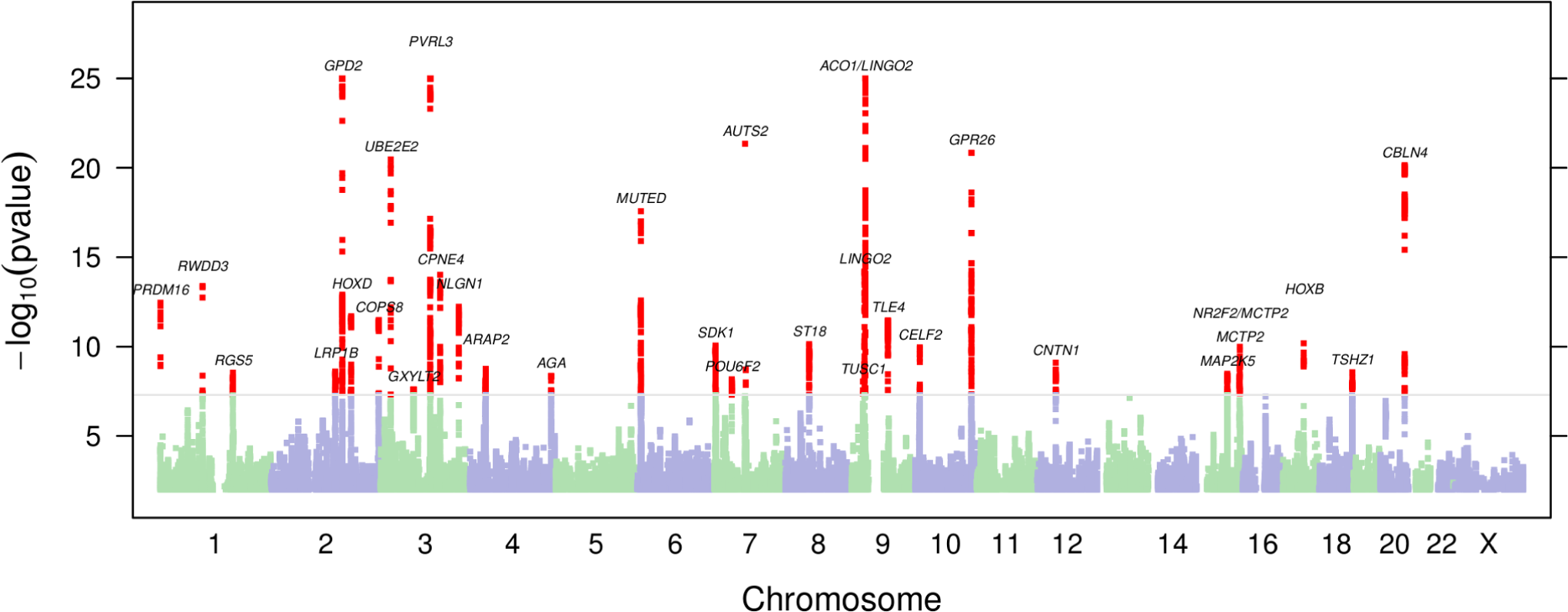
Manhattan plot. The 35 genome-wide significant regions are listed with the proposed candidate gene; regions that are close together share a label.

**Table 1:** Cohort statistics for motion sickness GWAS. Degree of motion sickness stratified by sex and age. Females and younger people tend to be more motion sick.

| Group | Total | Male | Female | $\leq 30$ | 31–45 | 46–60 | $\geq 61$ |
| --- | --- | --- | --- | --- | --- | --- | --- |
| Never | 40,042 | 25,137 | 14,905 | 5,510 | 10,707 | 10,592 | 13,233 |
| Occasionally | 24,902 | 11,855 | 13,047 | 4,597 | 8,451 | 6,459 | 5,395 |
| Sometimes | 6,723 | 3,067 | 3,656 | 1,175 | 2,015 | 1,759 | 1,774 |
| Frequently | 8,827 | 3,011 | 5,816 | 1,784 | 3,173 | 2,310 | 1,560 |
| Total | 80,494 | 43,070 | 37,424 | 13,066 | 24,346 | 21,120 | 21,962 |

Lead SNPs with p-values under 5 × 10^−8^ for motion sickness are shown in Table 2; 35 regions were significant (Figure **??**). We created a genetic propensity score based on the number of risk alleles for the 35 index SNPs. Individuals in the top five percent of the distribution (allele dosage of 40.25 or more risk alleles) had an average motion-sickness score 0.546 units higher than those in the bottom five percent (28.37 or fewer risk alleles). The top five percent had 6.37 times increased odds of being “frequently” motion sick as opposed to “never” motion sick as compared to the bottom five percent. The variance in motion sickness explained by the propensity score (which may be inflated as it was assessed in the discovery population) was 0.029.

**Table 2:**
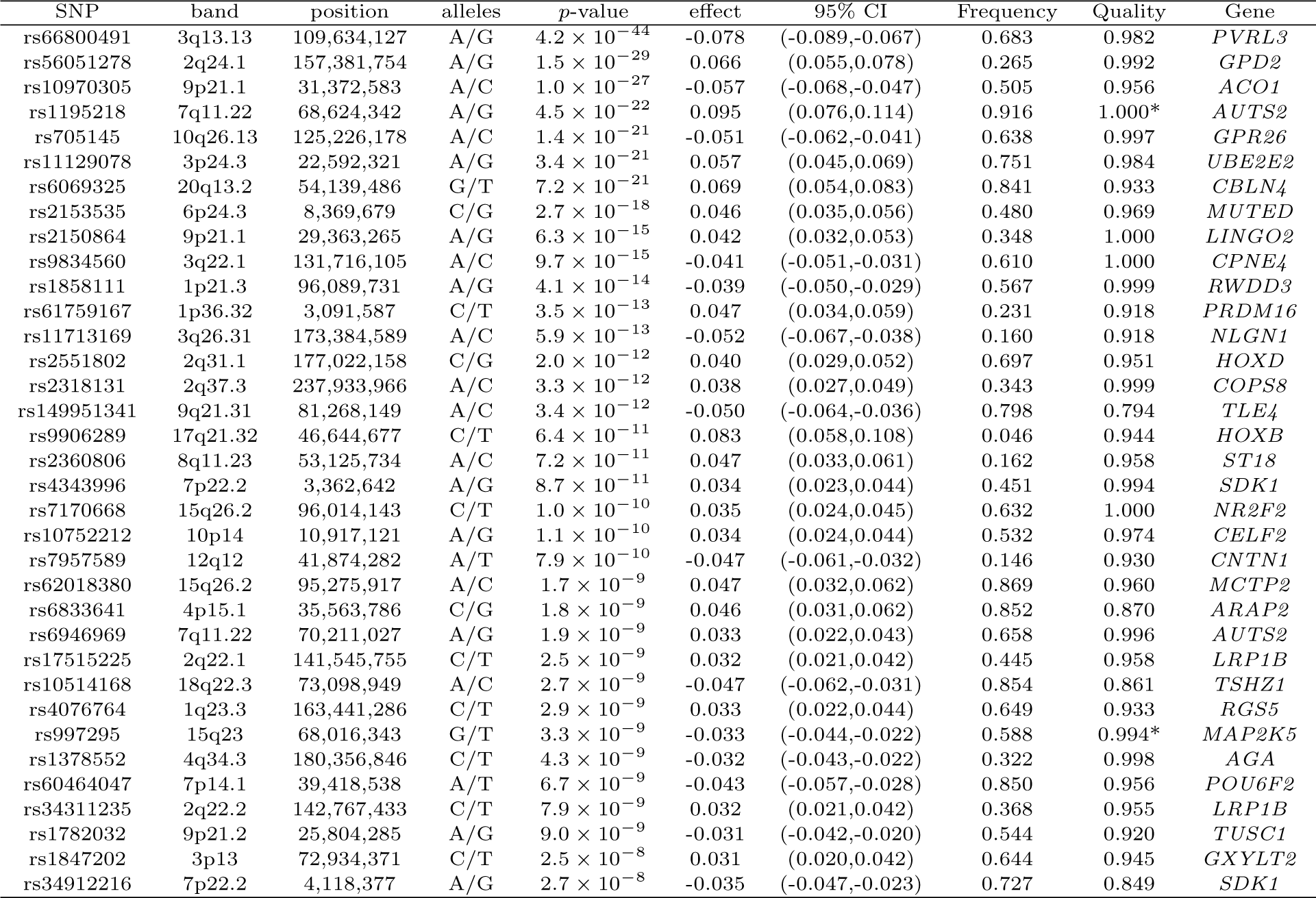
Genome-wide significant index SNPs. Alleles are reported in alphabetical order with respect to the positive strand of build 37 of the human genome. The effect is the change per copy of the second allele on a four point scale of increasing motion sickness. Frequency is the frequency of the alphabetically second allele in the cohort. Quality is imputation *r*^2^ for imputed SNPs, call rate for genotyped SNPs (those marked with a *). Gene is a proposed candidate gene in the region.

A few associated SNPs are in regions implicated in eye and ear development or balance. For example, our most significant association is with rs66800491 (*p* = 4.2 × 10^−44^), located roughly one Mb upstream of *PVRL3*, which encodes the cell adhesion protein Nectin-3. Loss of *PVRL3* expression in both humans and mice results in ocular defects [19]. The SNP rs10514168 (*p* = 2.7 × 10^−9^) is located down-stream of *TSHZ1*, a gene involved in inner ear development in the mouse [20]. Another association is with rs2153535 (*p* = 2.7 × 10^−18^), located upstream of *MUTED*, which is implicated in balance [21]. Three additional associated SNPs are near genes with major roles in early development: rs2551802 (*p* = 2 × 10^−12^) between *HOXD3* and *HOXD4*; rs9906289 (*p* = 6.4 × 10^−11^) in *HOXB3*; and rs149951341 (*p* = 3.4 × 10^−12^) near *TLE4*. The *HOXD* SNP is in LD (*r*^2^ ≈ 0.9) with rs2072590, which is associated with ovarian cancer [22].

Several other associated SNPs are located near genes involved in neurological processes including synapse development and function: rs11713169 (*p* = 5.9 × 10^−13^) in *NLGN1* encoding neuroligin; rs6069325 (*p* = 7.2 × 10^−21^) upstream of *CBLN4* encoding a member of the cerebellin precursor protein family; rs62018380 (*p* = 1.7 × 10^−9^) downstream of *MCTP2*, a gene involved in intercellular signal transduction and synapse function; rs7957589 in *PDZRN4* (*p* = 7.9 × 10^−10^) near *CNTN1* (contactin 1), which plays a role in axon guidance during neural development [23]; and two independent SNPs, rs4343996 and rs34912216 (*p* = 8.7 × 10^−11^ and 2.7 × 10^−8^, respectively) in *SDK1* encoding sidekick-1, a cell adhesion molecule that localizes to synapses. The SNP rs2150864 (*p* = 6.3 × 10^−15^) is located about 1.5 Mb upstream of *LINGO2*, a gene implicated in essential tremor [24]. Additional associated SNPs in or near genes in neurological pathways include: rs9834560 (*p* = 9.7 × 10^−15^) in *CPNE4* encoding copine-4, and two independent SNPS in or near *AUTS2* (rs1195218 and rs6946969 (*p* = 4.5 × 10^−22^ and 1.9 × 10^−9^, respectively).

Other associated SNPs are in regions involved in glucose and insulin homeostasis. For example, the second most significant association we found is with rs56051278 (*p* = 1.5 × 10^−29^) in *GPD2* that encodes glycerol-3-phosphate dehydrogenase 2, an enzyme implicated in glucose homeostasis. This SNP is in high LD (*r*^2^ ≈ 0.8) with rs2116665 (the non synonymous substitution H264R in *GPD2)* that was previously associated with free fatty acid and glycerol levels [25]. The SNP rs11129078 (*p* = 3.4 × 10^−21^) is located downstream of *UBE2E2*, which encodes a component of the ubiquitin-proteasome system. This system is implicated in the autophagy of pancreatic beta-cells that produce insulin and plays important roles in insulin homeostasis [26]. In addition, rs705145 (*p* = 1.4 × 10^−21^) is located just upstream of *GPR26*, encoding a G protein-coupled receptor. Mice with a deletion of the *GPR26* gene develop hyperphagia and diet-induced obesity, which leads to metabolic complications linked to obesity including glucose intolerance, hyperinsulemia and dyslipidemia [27]. The SNP rs4076764 (*p* = 2.9 × 10^−9^) is located upstream of *RGS5*, a regulator of G protein signaling. Loss of *RGS5* in the mouse is also associated with hyperphagia [28]. Finally, rs7170668 (*p* = 1 × 10^−10^) is located upstream of *NR2F2* encoding COUP-TFII (chicken ovalbumin upstream promoter transcription factor II), a protein with roles in glucose homeostasis and energy metabolism [29].

The remaining associated SNPs are in regions implicated in hypoxia (rs1858111 near *RWDD3*, *p* = 4.1 × 10^−14^); iron homeostasis (rs10970305 near *ACO1*, *p* = 1 × 10^−27^); brown adipose tissue (rs61759167 in *PRDM16*, *p* = 3.5 × 10^−13^); and other less characterized processes: rs2360806 (*p* = 7.2 × 10^−11^) in *ST18*, rs2318131 (*p* = 3.3 × 10^−12^) near *COPS8*, rs60464047 (*p* = 6.7 × 10^−9^) in *POU6F2*, rs10752212 (*p* = 1.1 × 10^−10^) near *CELF2*, rs6833641 (*p* = 1.8 × 10^−9^) near *ARAP2*, rs17515225 and rs34311235 (independently associated, *p* = 2.5 × 10^−9^ and 7.9 × 10^−9^, respectively) in *LRP1B*, rs1378552 (*p* = 4.3 × 10^−9^) in a gene desert on 4q34.3; and rs1782032 (*p* = 9 × 10^−9^) near *TUSC1*, and rs1847202 near *SHQ1* and *GXYLT2*. Finally, rs997295 in *MAP2K5* (*p* = 3.3 × 10^−9^) is in LD with rs2241423 (*r*^2^ ≈ 0.36), which is associated with BMI [30].

### Enrichment

Analysis of all regions with *p
* < 10^−5^ using GREAT [31] showed a significant enrichment in regions containing genes involved in fusion of atlas and occipital bones (FDR = 0.002) and abnormal arcus anterior morphology (FDR = 0.038) in mouse. The genes annotated with one or both of these processes were *HOXB*, *HOXD*, *TSHZ1*, and *RARB* regions (the SNP near *RARB* is rs2067120, *p* = 8.2 × 10^−6^).

### Phenotypic Study of Motion Sickness

We investigated comorbidities with motion sickness within the 23andMe database. Briefly, we looked at partial correlations between each of 695 different phenotypes and motion sickness, controlling for age, sex (where applicable), and 5 principal components. Table 3 shows selected large correlations. Some of the associated phenotypes are known symptoms of motion sickness (e.g., headache) or established comorbidities (migraines, vertigo, PONV, and morning sickness). In addition to PONV, other gastrointestinal (GI) phenotypes were also associated with motion sickness (e.g., irritable bowel syndrome (IBS); acid reflux; stomach upset with antidepressants, codeine and nonsteroidal anti-inflammatory drugs (NSAIDs); and indigestion with dairy products). Other associations include poor sleep, poor circulation, altitude sickness, hay fever, and neuroticism. Phenotypes associated with lower risk for motion sickness include a history of tobacco use, a good sense of direction, higher BMI, being single, and a better ability to handle stress.

**Table 3:** Selected partial correlations with motion sickness. Partial correlations are controlled for age, sex, and 5 PCs. *N* refers to the number of people with data for both motion sickness and the second trait. Traits are sorted by partial correlation.

| Phenotype | $\rho$ | $N$ |
| --- | --- | --- |
| Dizziness | 0.119 | 24606 |
| PONV | 0.117 | 25223 |
| Lightheaded during exercise | 0.114 | 23434 |
| Vomiting from codeine | 0.114 | 12176 |
| Altitude sickness | 0.111 | 41661 |
| Morning sickness | 0.108 | 13285 |
| Daytime sleepiness | 0.080 | 29384 |
| Indigestion due to dairy products | 0.071 | 29659 |
| Hay fever | 0.069 | 22242 |
| Headache after red wine | 0.066 | 42485 |
| Vertigo | 0.065 | 54934 |
| Back pain frequency | 0.065 | 28808 |
| Neuroticism | 0.063 | 38711 |
| Rosacea | 0.063 | 24635 |
| Irritable bowel syndrome (IBS) | 0.062 | 34348 |
| Mosquito bites itching more | 0.061 | 51292 |
| Greater perceived stress | 0.060 | 32486 |
| More colds last year | 0.060 | 34759 |
| Drowsiness from Benadryl | 0.055 | 19307 |
| Migraines | 0.055 | 72901 |
| Seasonal affective disorder | 0.051 | 32273 |
| GI pain from NSAIDs | 0.047 | 29846 |
| More sleep needed | 0.046 | 39115 |
| Nausea from antidepressants | 0.046 | 12079 |
| Dry mouth from antihistamines | 0.033 | 15826 |
| Acid reflux | 0.033 | 68998 |
| Being single | -0.033 | 65186 |
| Adventurous | -0.033 | 34532 |
| BMI | -0.034 | 75217 |
| Punctuality | -0.035 | 37486 |
| Drive through food frequency | -0.035 | 41467 |
| Good sense of direction | -0.039 | 46440 |
| Positive attitude towards self | -0.040 | 34082 |
| Pack years (cigarettes) | -0.046 | 66381 |
| Sound sleeper | -0.047 | 48339 |
| Ability to handle stress | -0.072 | 30965 |

### Genetic Correlations Between Motion Sickness and Related Phenotypes

We determined if any of the 35 SNPs associated with motion sickness were also associated with six correlated and clinically important phenotypes (PONV, migraines, hay fever, altitude sickness, morning sickness, and vertigo). Table 4 shows SNPs associated with these phenotypes with a (Bonferroni-corrected) *p*-value under 0.05/35 ≈ 0.0014 (under a more strinent threshold of 0.05/(35 × 6) ≈ 0.0002 only the first two are significant). One motion sickness associated SNP was significantly associated with migraines: rs61759167 in *PRDM16* (*p* = 1.1 × ^−6^). A previous study [32] reported an association between migraines and another SNP in *PRDM16*, rs2651899, which is in weak LD with rs61759167 (*r*^2^ ≈ 0.44). Three motion sickness-associated SNPs were also significantly associated with PONV: rs6833641 near *ARAP2*, rs1195218 near *AUTS2*, and rs6069325 near *CBLN4*. For all four examples, the higher risk allele for migraines or PONV is also the higher risk allele for motion sickness. We did not detect significant associations between motion sickness-associated SNPs and altitude sickness, hay fever, morning sickness, or vertigo. While this data suggests some shared etiology for motion sickness and PONV or migraines, it is difficult to assess whether or not this is due to shared causal SNPs.

### Sex-specific effects

Motion sickness is much more common in women than in men (Table 1) and several of our SNPs show much stronger effects in women than in men. The SNP rs66800491 has a 1.5x larger effect in women (− 0.097 versus −0.062) and rs1847202 has a 3x larger effect in women (0.048 versus 0.016) (both SNPs *p* < 0.05 for interaction, corrected for 35 tests). Overall 26 of the 35 SNPs have estimated larger effects in women than men (binomial *p* < 0.003; Table S1).

**Table 4:** Significant associations between motion sickness-associated SNPs and other phenotypes. *N* is the number of people with data for motion sickness and migraines or PONV.

| Phenotype | SNP | *N* | pvalue | effect | 95% CI |
| --- | --- | --- | --- | --- | --- |
| Migraine | rs61759167 | 72,901 | $1.1 \times 10^{-6}$ | 0.08 | (0.051, 0.119) |
| PONV | rs1195218 | 25,223 | 0.00012 | -0.14 | (-0.213, -0.069) |
| PONV | rs6069325 | 25,223 | 0.00079 | 0.09 | (0.038, 0.143) |
| PONV | rs6833641 | 25,223 | 0.00101 | 0.09 | (0.037, 0.148) |

## Discussion

Here we report 35 novel, genome-wide significant associations for motion sickness (Table 2). Genes in regions associated with motion sickness appear to play roles in eye and ear development, balance and other neurological processes, and glucose homeostasis. Two of the genome-wide significant regions contain hypoxia-inducible genes. We also provide evidence that motion sickness is phenotypically associated with numerous conditions and traits (Table 3).

Since motion sickness is thought to stem from the brain receiving contradictory signals from the inner ear versus the eye (e.g., the inner ear signals “moving” while the eye signals “stationary”), it is interesting that a region implicated in eye development (rs56100358 near *PVRL3)* is our most significant association. Chromosomal rearrangements that lead to loss of *PVRL3* expression have been associated with ocular defects in humans and the *PVRL3* knockout mouse exhibits lens and other vision problems [19]. The associations with regions involved in the inner ear (rs12111385 near *MUTED* and rs1435985 near *TSHZ1)* are also interesting since disturbances in the vestibular system of the inner ear, which senses motion and body position and influences balance, are thought to play a central role in motion sickness. It has been suggested that the mouse homolog of *MUTED* controls the synthesis of otoliths of the vestibular labyrinth of the inner ear [21]. Otoliths are sensitive to gravity and linear acceleration and play a role in balance. Mutations in *TSHZ1* and deletions in the 18q22.3 region that includes *TSHZ1* are associated with congenital aural atresia (CAA) [33, 34], a spectrum of ear deformities that involve malformation of the external auditory canal (EAC). More generally, our enrichment analysis suggests that genes involved in certain aspects of cranial developmental may play an important role in motion sickness. Associations with SNPs in or near genes involved in synapse formation and function (*NLGN1*, *CBLN4*, *MCTP2*, *PDZRN4*, *CNTN1*, and *SDK1)* and other neurological pathways (*LINGO2*, *CPNE4*, *AUTS2)* point to the importance of the brain in motion sickness.

Five associated SNPs are in or near genes implicated in glucose and insulin homeostasis or BMI. Although these SNPs are not in linkage disequilibrium (LD) with SNPs reported in GWAS of type 2 diabetes (T2D) [35–40], rs56051278 is in high LD (*r*^2^ ≈ 0.8) with rs2116665, a nonsynonymous substitution (H264R) in the *GPD2* gene. H264R has been associated with increased plasma glycerol and free fatty acid concentrations in a French Canadian population [25]. Increased free fatty acid levels are indicative of glucose intolerance and hyperinsulinemia. Although it is unclear why genes involved in glucose and insulin regulation might also play a role in motion sickness, one study suggested that hyperglycemia may be related to the gastrointestinal symptoms of motion sickness [15]. In this study, individuals who experienced motion-induced nausea and vomiting had lower levels of insulin than people who did not experience gastrointestinal symptoms. The study further suggested that stable glucose levels might help to relieve motion-induced gastrointestinal upset.

At least two of our associated SNPs are near hypoxia-inducible genes: *RGS5* and *RWDD3* (encoding the RSUME protein). RSUME promotes the activity of hypoxia-inducible factor 1 (HIF-1alpha), a master regulator of the hypoxic response [41]; *RGS5* is an apoptotic stimulator induced by hypoxia in endothelial cells [42]. This data suggests a potential relationship between motion sickness and hypoxia. Motion sickness might lead to hypoxia or individuals predisposed to hypoxia might also be more susceptible to motion sickness. Both possibilities are intriguing since our phenotypic analysis suggested an association between motion sickness and altitude sickness, which occurs when individuals become hypoxic at higher altitudes (Table 3).

Among the regional association plots (Supplement), one SNP in particular stands out: rs1195218 near *AUTS2*. This genotyped SNP has a *p*-value under 10^−20^ and no other SNPs in the region have *p* < 10^−6^. This lack of signal from LD is not terribly surprising, as none of the three proxy SNPs (*r*^2^ > 0.2) for this SNP in 1000 Genomes pass imputation quality control in our data. As the clusters for this SNP look excellent and the call rate is 99.98%, we believe this is a true signal.

Certain phenotypic associations are interesting given what is known about motion sickness. PONV is an established comorbidity of motion sickness [43] and is thought to stem from the anaesthetics that are administered for surgery. It may not, therefore, be surprising that motion sickness was also associated with vomiting and/or nausea with use of codeine, antidepressants and nonsteroidal anti-inflammatories (NSAIDs). Additional gastrointestinal phenotypes (e.g., IBS, acid reflux, and indigestion with dairy products) as well as other drug-related phenotypes like being drowsy when taking Benadryl and feeling jittery when taking Sudafed were also associated with motion sickness. Interestingly, our findings suggest shared genetic susceptibility for both motion sickness and PONV (Table 4).

Some phenotypic associations might provide clues about the etiology of motion sickness (e.g., poor circulation and becoming light headed with exercise) or they might suggest simple remedies for motion sickness such as improving sleep quality. A number of associated phenotypes were related to personality (e.g., neuroticism) or behavior (e.g., smoking). We note, however, that it is difficult to assess causality for these phenotype-phenotype associations. For example, does being a sound sleeper make one less susceptible to being motion sick, or vice versa, or are both related to a third condition? The validity of these novel phenotypic findings is bolstered by the fact that we also detected associations with known symptoms (dizziness and headache) and established comorbidities (PONV, migraines, vertigo and morning sickness) of motion sickness. In some cases we even identified shared genetic factors for motion sickness and related comorbidities (e.g., PONV and migraines). Some of the correlated phenotypes are not established comorbidities or symptoms of motion sickness, however, and do not have an obvious biological relationship to motion sickness.

Our web-based method of capturing phenotypic information allows us to build a very large cohort (e.g., 80,494 individuals in our study), but we may not have obtained a complete picture of an individual’s susceptibility to motion sickness. For finding SNPs, the gain in power from having a large sample more than makes up for the reduction in power due to possible misclassification. An additional potential limitation is that we only surveyed individuals about car sickness; future studies should investigate these SNPs in populations phenotyped for other forms of motion sickness. An advantage of our web-based phenotypic collection method is that we can easily investigate whether seemingly related traits have shared underlying genetics. We identified four SNPs simultaneously associated with motion sickness plus PONV or migraines. These findings may provide clues into the etiology of all three conditions and may point to overlapping risk factors or treatments.

## Methods

### Human Subjects

All participants were drawn from the customer base of 23andMe, Inc., a consumer genetics company. This cohort has been described in detail previously [44,45]. Participants provided informed consent and participated in the research online, under a protocol approved by the external AAHRPP-accredited IRB, Ethical & Independent Review Services (E&I Review).

### Phenotype collection

Participants invited to fill out web-based questionnaires, which included four questions about motion sickness during road travel, whenever they logged into their 23andMe accounts. The responses to each question were translated into a motion sickness score of 0 (never), 1 (occasionally), 2 (sometimes), or 3 (frequently). Responses of “I’m not sure” or “Don’t know” were excluded from the analysis. The questions were prioritized (1–4) and a participant’s final score was based on the question they answered that had the highest priority. The questions were:

1. “How often do you experience motion sickness while in a car? (Never / Occasionally / Sometimes / Frequently / I’m not sure)”
2. “Have you experienced motion sickness while riding in a car (car sickness)? (Yes, I do now frequently / Yes, I did frequently, but only as a child / 3. Yes, occasionally / No / Don’t know)”
3. “As a child, how often did you experience motion sickness while in a car? (Never / Occasionally / Sometimes / Frequently / I’m not sure)”
4. “Can you read in a moving car without becoming nauseated? (Never / Sometimes / Always / I’m not sure)”

Responses to the final question were scored as Never = 3, Sometimes = 1, and Always = 0.

### Genotyping and imputation

Participants were genotyped and additional SNP genotypes were imputed against the August 2010 release of the 1000 Genomes data as described previously [46]. Briefly, they were genotyped on at least one of three genotyping platforms, two based on the Illumina HumanHap550+ BeadChip, the third based on the Illumina Human OmniExpress+ BeadChip. The platforms included assays for 586,916, 584,942, and 1,008,948 SNPs, respectively. Genotypes for a total of 11,914,767 SNPs were imputed in batches of roughly 10,000 individuals, grouped by genotyping platform. Imputation was performed as in [46]. Prior to the imputation we discarded genotyped SNPs that were not present in the imputation panel. For the GWAS, we added such SNPs back (if they passed quality control), for a total of 7,428,049 SNPs (7,378,897 imputed and 49,152 genotyped). To filter SNPs whose imputation results had changed over time, we performed an ANOVA test for frequency differences across batches. The quality control criteria for imputed SNPs were batch effects *p*-value at least 10^−50^, average *r*^2^ across batches of at least 0.5, and minimum *r*^2^ across batches of at least 0.3. For genotyped SNPs, we required a MAF of at least 0.001, a Hardy-Weinburg *p*-value of at least 10^−20^, and a call rate at least 0.9.

### Statistical analysis

In order to minimize population substructure while maximizing statistical power, the study was limited to individuals with European ancestry. Ancestry was inferred from the genome-wide genotype data and principal component analysis was performed as in [44, 47]. The cohort was filtered by relatedness to remove participants at a first cousin or closer relationship. More precisely, no two participants shared more than 700 cM of DNA identical by descent (IBD; approximately the lower end of sharing between a pair of first cousins). IBD was calculated using the methods described in [48]. The genomic control inflation factor was 1.156; all p-values are adjusted for this inflation factor.

The GWAS was performed using likelihood ratio tests for the linear regression

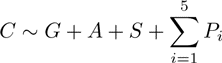

of carsickness on genotype, age, sex and 5 principal components of genetic ancestry. Genotypes were coded as a dosage from 0–2 (counting the estimated number of minor alleles present for imputed SNPs) or as a count 0, 1, or 2 (also number of minor alleles, for genotyped SNPs). Significant SNPs were grouped into regions with at least 500 kb between pairs of significant SNPs; the SNP with the lowest *p*-value in each region was chosen to be the index SNP. As some of the regions were under 1Mb apart, a joint regression with all index SNPs was run to make sure that they all represented independent signals.

Partial correlation between carsickness *C* and a phenotype *Y* were computed by computing the correlation between the residuals produced by regressing both *C* and *Y* on age, sex, and five principal components, using linear regression even if *Y* was a binary trait. We did not attempt to quantify the significance of these regressions nor any causality.

Tests of SNPs associated with motion sickness against other correlated traits were done using logistic or linear regressions as appropriate with the same covariates as in the GWAS (except for morning sickness, which dropped sex). The phenotypes studied (PONV, migraines, hay fever, altitude sickness, morning sickness, and vertigo) were all case control except for morning sickness, which was scored on a 5 point scale: None; Mild (occasional bouts of queasiness or nausea, did not require treatment); Moderate (nausea and some vomiting, but did not require treatment); Severe (Severe nausea and vomiting that required treatment); Very severe (requiring hospitalization and intravenous fluid (IV) therapy).

Enrichment analysis using GREAT was conducted on all regions with an index SNP with *p* < 10^−5^, where regions were enforced to be 500 kb apart. Windows of 500 kb on eather side of each index SNP were uploaded into GREAT using default settings.

## Acknowledgements

We thank the customers of 23andMe who answered surveys and participated in this research. We also thank all the employees of 23andMe, who together have made this research possible.

## Supplement

**Table S1:** Sex-specific effects for significant SNPs. Estimated effect sizes for females, males, and their ratio. *p*-values are for a non-zero interaction term between sex and genotype. SNPs are sorted by increasing *p*-value.

| SNP | β-female | β-male | ratio | $p$-value |
| --- | --- | --- | --- | --- |
| rs66800491 | -0.097 | -0.062 | 1.549 | 0.001 |
| rs1847202 | 0.048 | 0.016 | 3.108 | 0.001 |
| rs6069325 | 0.089 | 0.050 | 1.772 | 0.004 |
| rs149951341 | -0.070 | -0.033 | 2.098 | 0.006 |
| rs61759167 | 0.060 | 0.035 | 1.735 | 0.032 |
| rs1195218 | 0.115 | 0.077 | 1.505 | 0.037 |
| rs1378552 | -0.044 | -0.023 | 1.935 | 0.039 |
| rs11129078 | 0.068 | 0.047 | 1.438 | 0.065 |
| rs2150864 | 0.051 | 0.035 | 1.472 | 0.104 |
| rs997295 | -0.042 | -0.025 | 1.671 | 0.106 |
| rs10970305 | -0.065 | -0.051 | 1.296 | 0.127 |
| rs34311235 | 0.040 | 0.024 | 1.630 | 0.128 |
| rs10752212 | 0.041 | 0.027 | 1.514 | 0.150 |
| rs62018380 | 0.057 | 0.038 | 1.519 | 0.178 |
| rs34912216 | -0.043 | -0.028 | 1.546 | 0.194 |
| rs10514168 | -0.056 | -0.038 | 1.475 | 0.215 |
| rs1858111 | -0.034 | -0.044 | 0.760 | 0.273 |
| rs9906289 | 0.096 | 0.071 | 1.354 | 0.288 |
| rs6946969 | 0.027 | 0.037 | 0.730 | 0.319 |
| rs2551802 | 0.046 | 0.036 | 1.292 | 0.330 |
| rs56051278 | 0.071 | 0.062 | 1.161 | 0.363 |
| rs6833641 | 0.040 | 0.052 | 0.767 | 0.400 |
| rs11713169 | -0.058 | -0.047 | 1.233 | 0.415 |
| rs1782032 | -0.027 | -0.035 | 0.774 | 0.434 |
| rs7170668 | 0.038 | 0.031 | 1.224 | 0.482 |
| rs60464047 | -0.047 | -0.040 | 1.182 | 0.601 |
| rs705145 | -0.049 | -0.053 | 0.910 | 0.630 |
| rs2153535 | 0.043 | 0.048 | 0.903 | 0.633 |
| rs9834560 | -0.043 | -0.039 | 1.098 | 0.696 |
| rs2318131 | 0.040 | 0.036 | 1.106 | 0.706 |
| rs7957589 | -0.044 | -0.049 | 0.917 | 0.776 |
| rs4076764 | 0.034 | 0.032 | 1.067 | 0.836 |
| rs4343996 | 0.033 | 0.034 | 0.949 | 0.856 |
| rs2360806 | 0.046 | 0.048 | 0.976 | 0.931 |
| rs17515225 | 0.032 | 0.031 | 1.018 | 0.954 |

**Figure S1:**
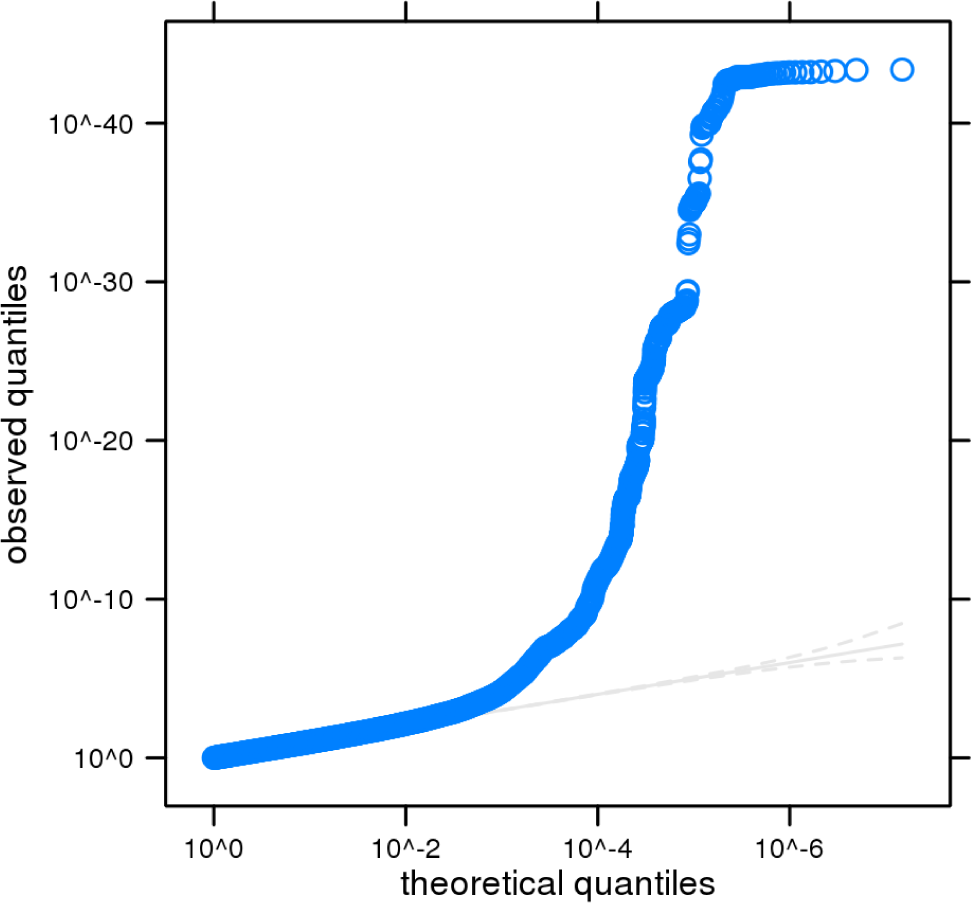
Quantile-quantile plot. Observed versus expected *p*-values under the null.

**Figure S2:**
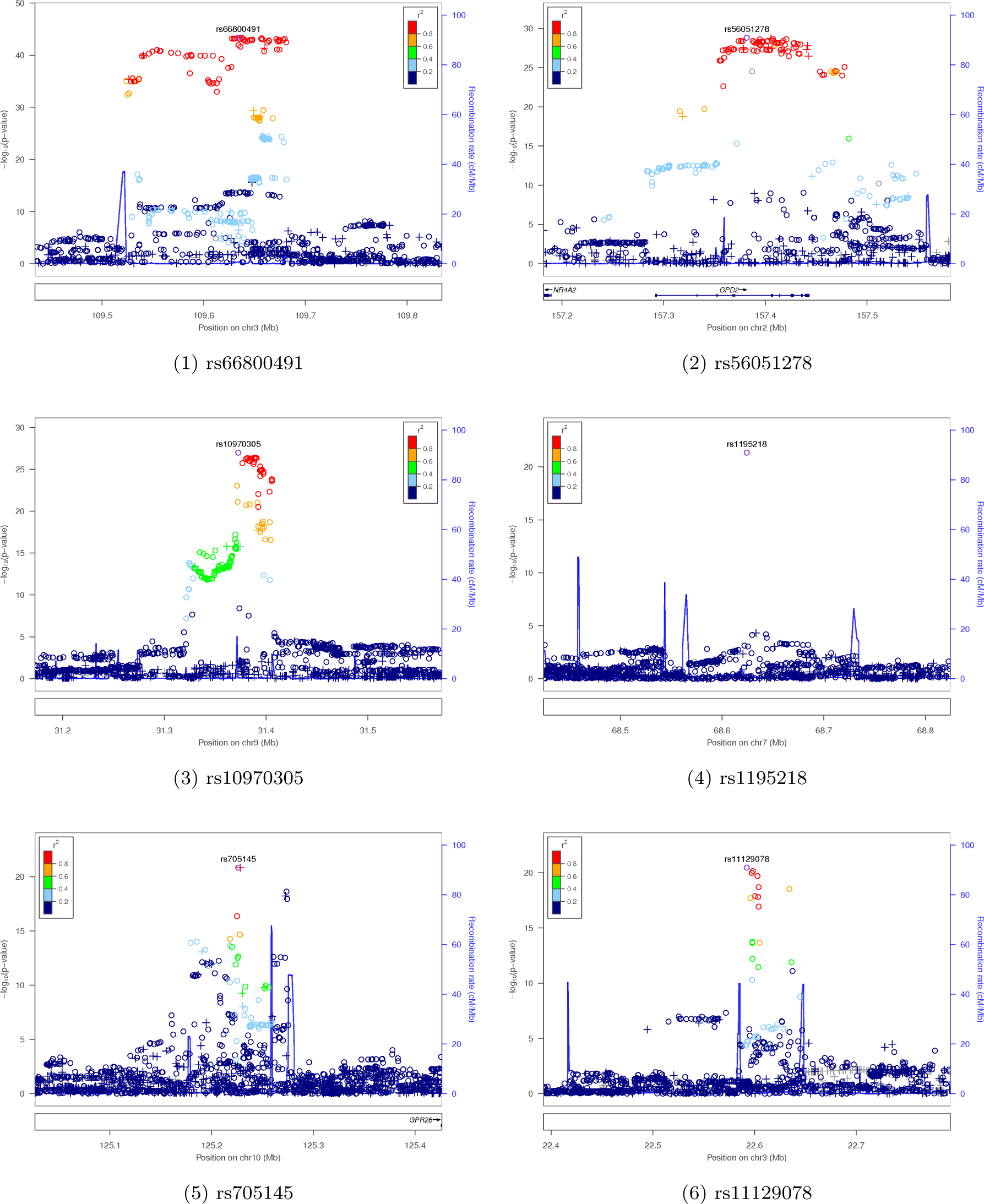
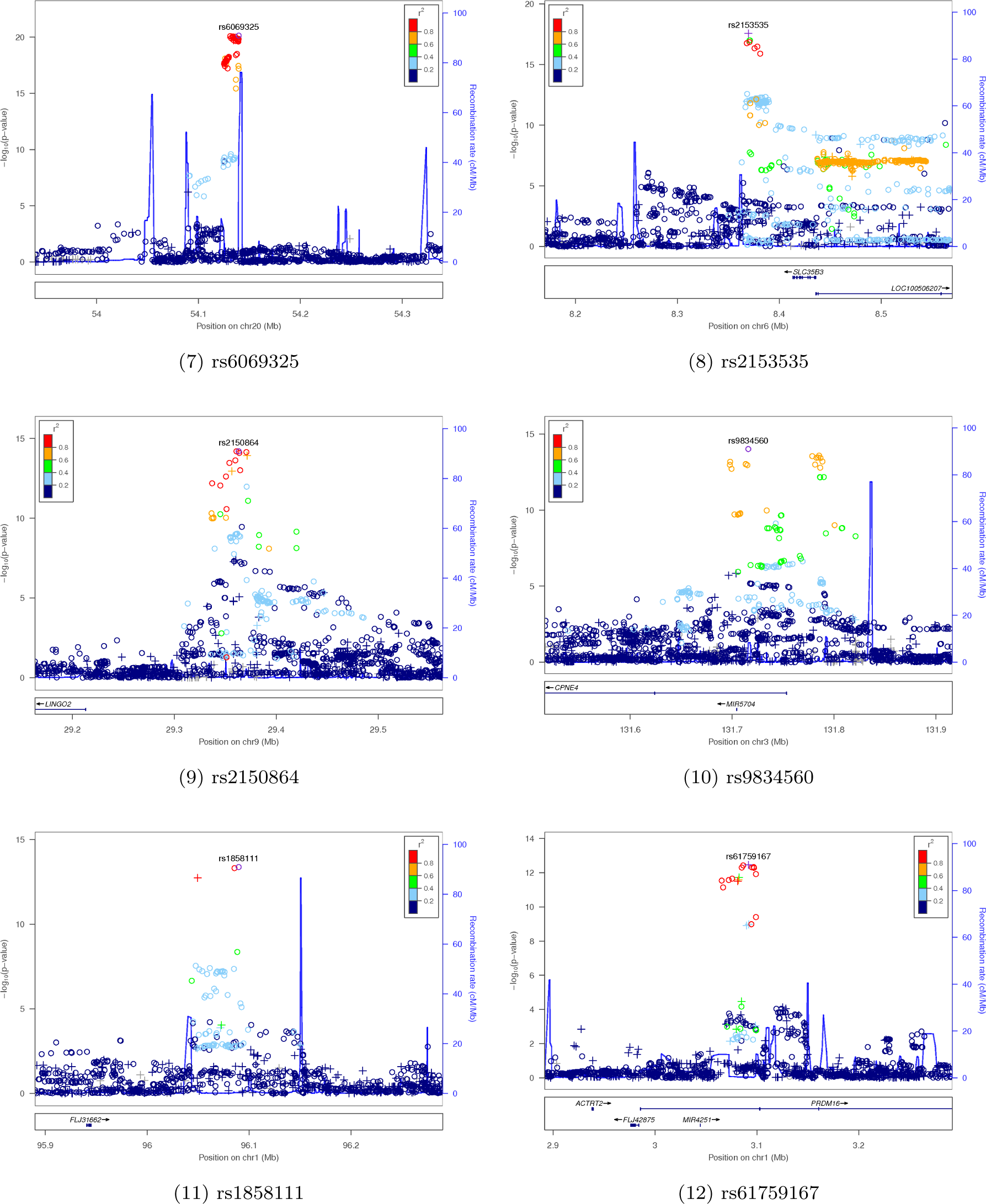
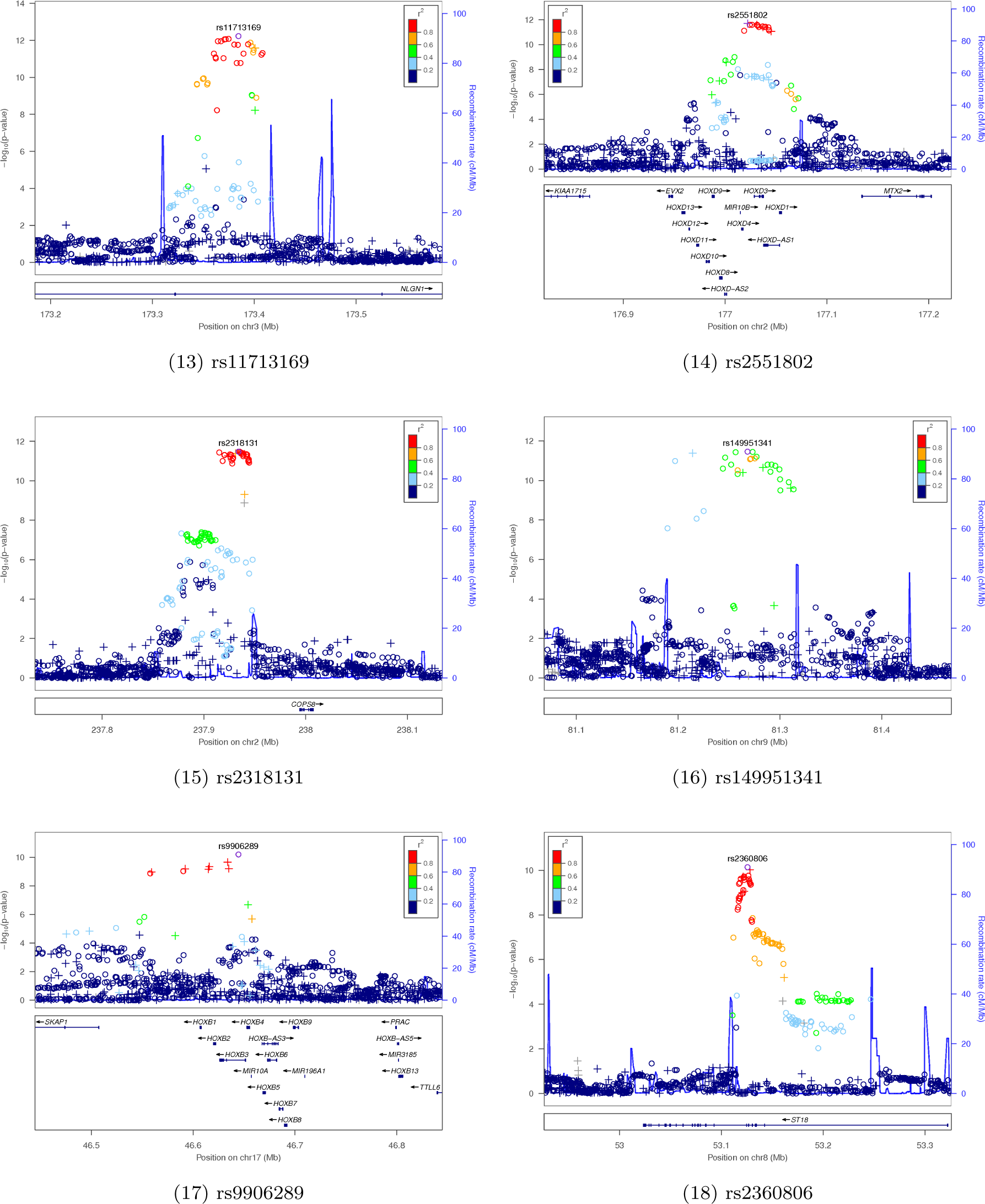
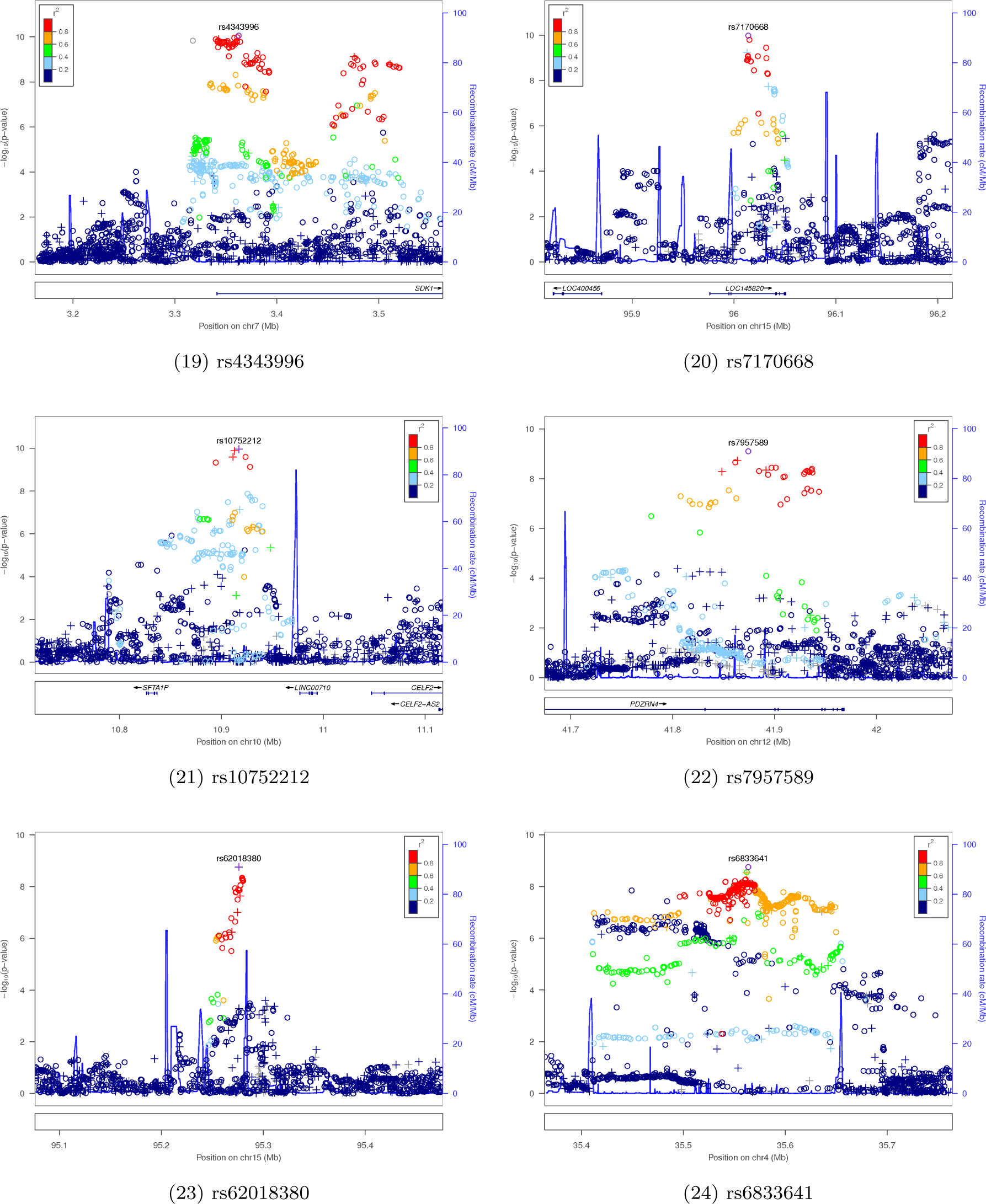
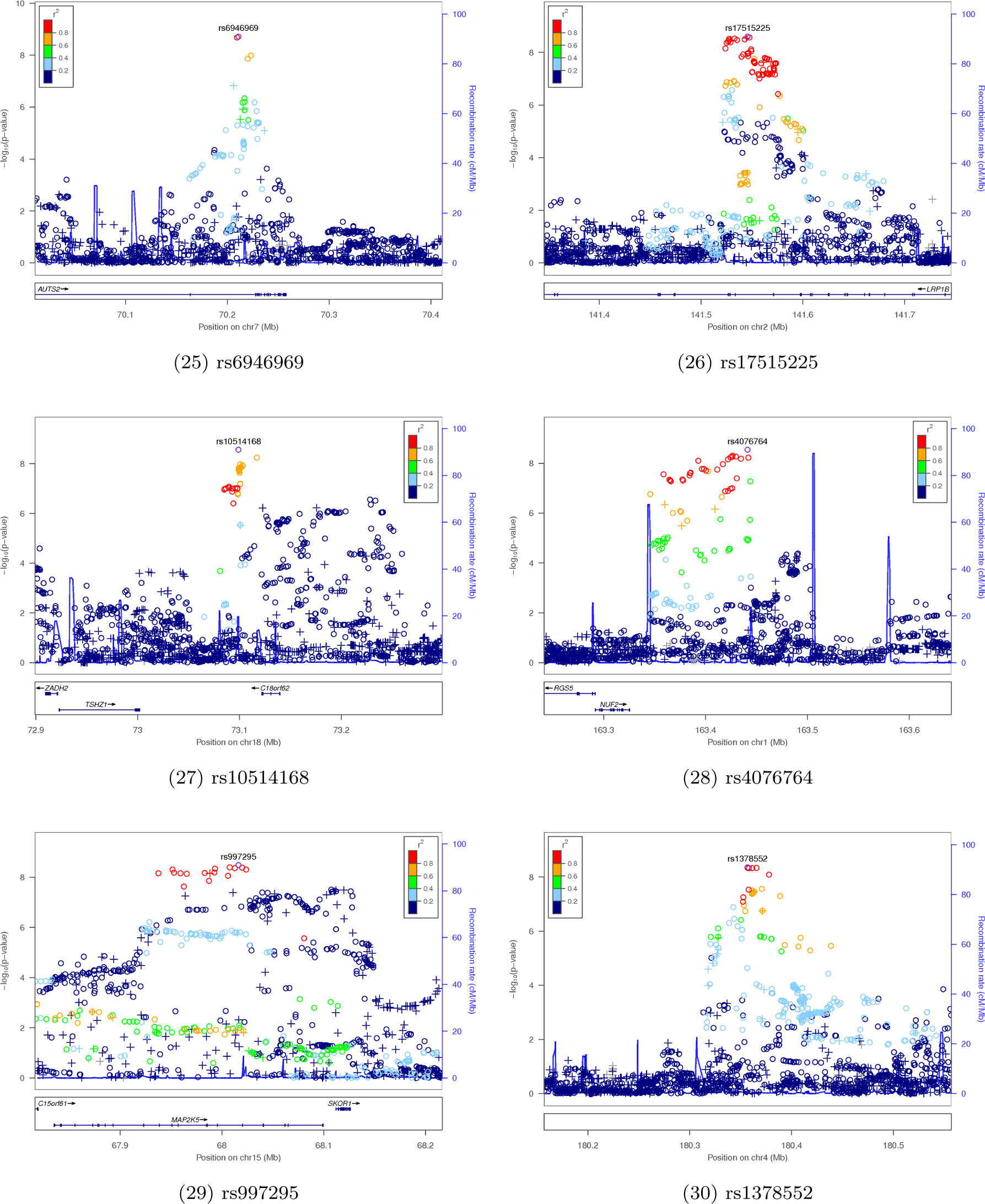
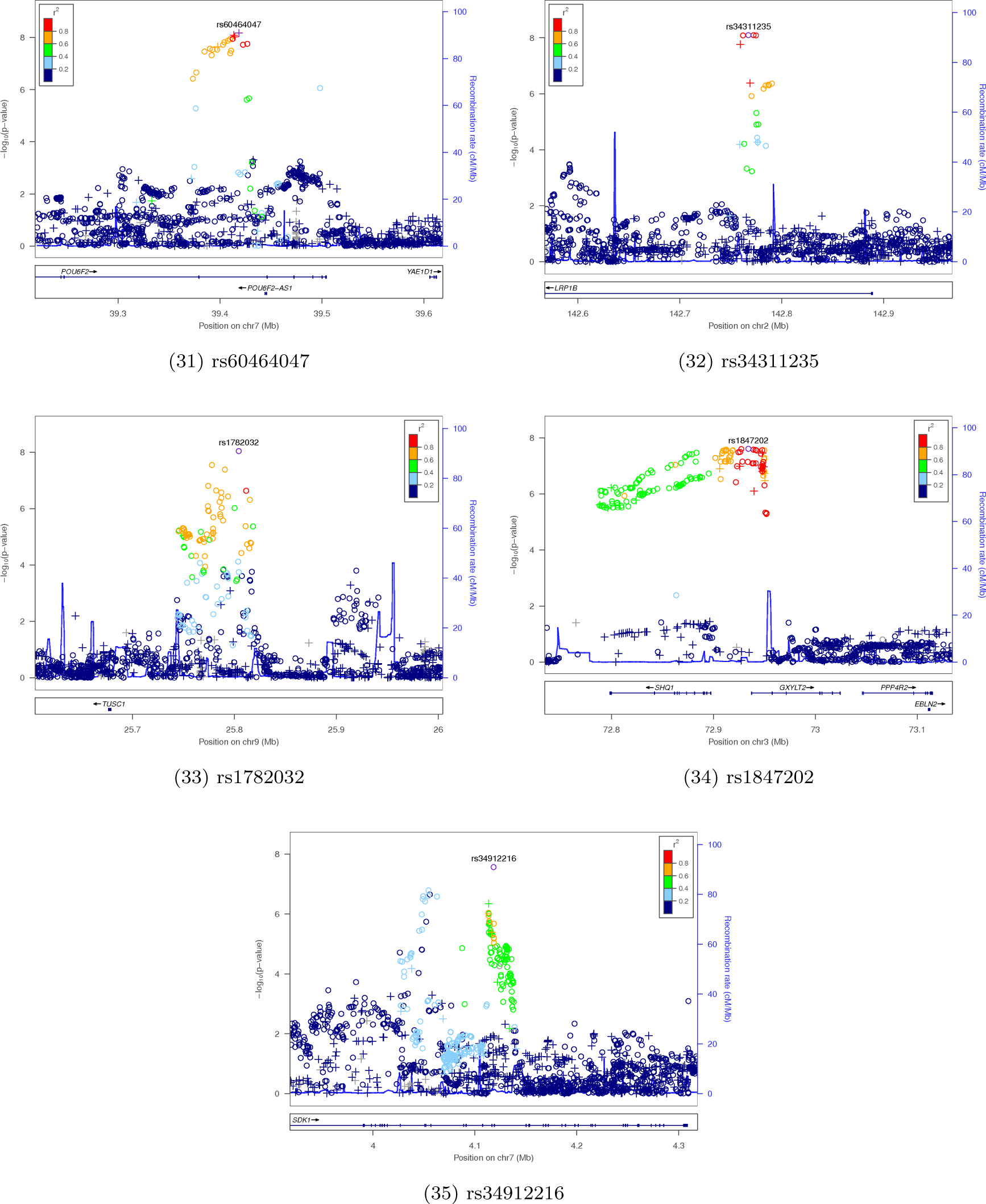
Region plots for genome-wide significant regions. SNPs are colored by *r*^2^ with the index SNP (which is labeled). Circles are imputed SNPs, plus signs are genotyped SNPs.

## References

[1] Covanis A (2006) Panayiotopoulos syndrome: a benign childhood autonomic epilepsy frequently imitating encephalitis, syncope, migraine, sleep disorder, or gastroenteritis. Pediatrics 118: e1237–1243.

[2] Yang SN, Schlieski T, Selmins B, Cooper SC, Doherty RA, et al. (2012) Stereoscopic viewing and reported perceived immersion and symptoms. Optom Vis Sci 89: 1068–1080.

[3] Murdin L, Golding J, Bronstein A (2011) Managing motion sickness. BMJ 343: d7430.

[4] Cuomo-Granston A, Drummond PD (2010) Migraine and motion sickness: what is the link? Prog Neurobiol 91: 300–312.

[5] Sherman CR (2002) Motion sickness: review of causes and preventive strategies. J Travel Med 9: 251–256.

[6] Lentz JM, Collins WE (1977) Motion sickness susceptibility and related behavioral characteristics in men and women. Aviat Space Environ Med 48: 316–322.

[7] Lindseth G, Lindseth PD (1995) The relationship of diet to airsickness. Aviat Space Environ Med 66: 537–541.

[8] Turner M, Griffin MJ (1999) Motion sickness in public road transport: passenger behavior and susceptibility. Ergonomics 42: 444–461.

[9] Lawther A, Griffin MJ (1988) A survey of the occurrence of motion sickness amongst passengers at sea. Aviat Space Environ Med 59: 399–406.

[10] Stern RM, Hu S, Uijtdehaage SH, Muth ER, Xu LH, et al. (1996) Asian hypersusceptibility to motion sickness. Hum Hered 46: 7–14.

[11] Klosterhalfen S, Kellermann S, Pan F, Stockhorst U, Hall G, et al. (2005) Effects of ethnicity and gender on motion sickness susceptibility. Aviat Space Environ Med 76: 1051–1057.

[12] Yu Y, Chung HC, Hemingway L, Stoffregen TA (2013) Standing body sway in women with and without morning sickness in pregnancy. Gait Posture 37: 103–107.

[13] Meissner K, Enck P, Muth ER, Kellermann S, Klosterhalfen S (2009) Cortisol levels predict motion sickness tolerance in women but not in men. Physiol Behav 97: 102–106.

[14] Matchock RL, Levine ME, Gianaros PJ, Stern RM (2008) Susceptibility to nausea and motion sickness as a function of the menstrual cycle. Womens Health Issues 18: 328–335.

[15] Mo FF, Qin HH, Wang XL, Shen ZL, Xu Z, et al. (2012) Acute hyperglycemia is related to gastrointestinal symptoms in motion sickness: an experimental study. Physiol Behav 105: 394–401.

[16] Kohl RL (1985) Endocrine correlates of susceptibility to motion sickness. Aviat Space Environ Med 56: 1158–1165.

[17] Muth ER (2006) Motion and space sickness: intestinal and autonomic correlates. Auton Neurosci 129: 58–66.

[18] Reavley CM, Golding JF, Cherkas LF, Spector TD, MacGregor AJ (2006) Genetic influences on motion sickness susceptibility in adult women: a classical twin study. Aviat Space Environ Med 77: 1148–1152.

[19] Lachke SA, Higgins AW, Inagaki M, Saadi I, Xi Q, et al. (2012) The cell adhesion gene PVRL3 is associated with congenital ocular defects. Hum Genet 131: 235–250.

[20] Core N, Caubit X, Metchat A, Boned A, Djabali M, et al. (2007) Tshz1 is required for axial skeleton, soft palate and middle ear development in mice. Dev Biol 308: 407–420.

[21] Zhang Q, Li W, Novak EK, Karim A, Mishra VS, et al. (2002) The gene for the muted (mu) mouse, a model for Hermansky-Pudlak syndrome, defines a novel protein which regulates vesicle trafficking. Hum Mol Genet 11: 697–706.

[22] Goode EL, Chenevix-Trench G, Song H, Ramus SJ, Notaridou M, et al. (2010) A genome-wide association study identifies susceptibility loci for ovarian cancer at 2q31 and 8q24. Nat Genet 42: 874–879.

[23] Shimoda Y, Watanabe K (2009) Contactins: emerging key roles in the development and function of the nervous system. Cell Adh Migr 3: 64–70.

[24] Jasinska-Myga B, Wider C (2012) Genetics of essential tremor. Parkinsonism Relat Disord 18 Suppl 1: S138–139.

[25] St-Pierre J, Vohl MC, Brisson D, Perron P, Despres JP, et al. (2001) A sequence variation in the mitochondrial glycerol-3-phosphate dehydrogenase gene is associated with increased lasma glycerol and free fatty acid concentrations among French Canadians. Mol Genet Metab 72: 209–217.

[26] Hartley T, Brumell J, Volchuk A (2009) Emerging roles for the ubiquitin-proteasome system and autophagy in pancreatic beta-cells. Am J Physiol Endocrinol Metab 296: 1–10.

[27] Chen D, Liu X, Zhang W, Shi Y (2012) Targeted inactivation of GPR26 leads to hyperphagia and adiposity by activating AMPK in the hypothalamus. PLOS ONE 7: e40764.

[28] Deng W, Wang X, Xiao J, Chen K, Zhou H, et al. (2012) Loss of regulator of G protein signaling 5 exacerbates obesity, hepatic steatosis, inflammation and insulin resistance. PLOS ONE 7: e30256.

[29] Li L, Xie X, Qin J, Jeha GS, Saha PK, et al. (2009) The nuclear orphan receptor COUPTFII plays an essential role in adipogenesis, glucose homeostasis, and energy metabolism. Cell Metab 9: 77–87.

[30] Speliotes EK, Willer CJ, Berndt SI, Monda KL, Thorleifsson G, et al. (2010) Association analyses of 249,796 individuals reveal 18 new loci associated with body mass index. Nat Genet 42: 937–948.

[31] McLean CY, Bristor D, Hiller M, Clarke SL, Schaar BT, et al. (2010) GREAT improves functional interpretation of cis-regulatory regions. Nat Biotechnol 28: 495–501.

[32] Chasman DI, Schurks M, Anttila V, de Vries B, Schminke U, et al. (2011) Genome-wide association study reveals three susceptibility loci for common migraine in the general population. Nat Genet 43: 695–698.

[33] Dostal A, Nemeckova J, Gaillyova R, Vranova V, Zezulkova D, et al. (2006) Identification of 2.3-Mb gene locus for congenital aural atresia in 18q22.3 deletion: a case report analyzed by comparative genomic hybridization. Otol Neurotol 27: 427–432.

[34] Feenstra I, Vissers LE, Pennings RJ, Nillessen W, Pfundt R, et al. (2011) Disruption of teashirt zinc finger homeobox1 is associated with congenital aural atresia in humans. Am J Hum Genet 89: 813–819.

[35] Saxena R, Voight BF, Lyssenko V, Burtt NP, de Bakker PI, et al. (2007) Genome-wide association analysis identifies loci for type 2 diabetes and triglyceride levels. Science 316: 1331–1336.

[36] Burton PR, Clayton DG, Cardon LR, Craddock N, Deloukas P, et al. (2007) Genome-wide association study of 14,000 cases of seven common diseases and 3,000 shared controls. Nature 447: 661–678.

[37] Zeggini E, Weedon MN, Lindgren CM, Frayling TM, Elliott KS, et al. (2007) Replication of genome-wide association signals in UK samples reveals risk loci for type 2 diabetes. Science 316: 1336–1341.

[38] Sladek R, Rocheleau G, Rung J, Dina C, Shen L, et al. (2007) A genome-wide association study identifies novel risk loci for type 2 diabetes. Nature 445: 881–885.

[39] Steinthorsdottir V, Thorleifsson G, Reynisdottir I, Benediktsson R, Jonsdottir T, et al. (2007) A variant in CDKAL1 influences insulin response and risk of type 2 diabetes. Nat Genet 39: 770– 775.

[40] Scott LJ, Mohlke KL, Bonnycastle LL, Willer CJ, Li Y, et al. (2007) A genome-wide association study of type 2 diabetes in Finns detects multiple susceptibility variants. Science 316: 1341–1345.

[41] Carbia-Nagashima A, Gerez J, Perez-Castro C, Paez-Pereda M, Silberstein S, et al. (2007) RSUME, a small RWD-containing protein, enhances SUMO conjugation and stabilizes HIF-1alpha during hypoxia. Cell 131: 309–323.

[42] Jin Y, An X, Ye Z, Cully B, Wu J, et al. (2009) RGS5, a hypoxia-inducible apoptotic stimulator in endothelial cells. J Biol Chem 284: 23436–23443.

[43] Apfel CC, Heidrich FM, Jukar-Rao S, Jalota L, Hornuss C, et al. (2012) Evidence-based analysis of risk factors for postoperative nausea and vomiting. Br J Anaesth 109: 742–753.

[44] Eriksson N, Macpherson JM, Tung JY, Hon LS, Naughton B, et al. (2010) Web-based, participant-driven studies yield novel genetic associations for common traits. PLOS Genet 6: e1000993.

[45] Tung JY, Do CB, Hinds DA, Kiefer AK, Macpherson JM, et al. (2011) Efficient replication of over 180 genetic associations with self-reported medical data. PLOS ONE 6: e23473.

[46] Eriksson N, Benton GM, Do CB, Kiefer AK, Mountain JL, et al. (2012) Genetic variants associated with breast size also influence breast cancer risk. BMC Med Genet 13: 53.

[47] Eriksson N, Tung JY, Kiefer AK, Hinds DA, Francke U, et al. (2012) Novel associations for hypothyroidism include known autoimmune risk loci. PLOS ONE 7: e34442.

[48] Henn BM, Hon L, Macpherson JM, Eriksson N, Saxonov S, et al. (2012) Cryptic distant relatives are common in both isolated and cosmopolitan genetic samples. PLOS ONE 7: e34267.

